# Oxytocin modulates the intrinsic dynamics between attention-related large scale networks

**DOI:** 10.1101/371971

**Authors:** Fei Xin, Feng Zhou, Xinqi Zhou, Xiaole Ma, Yayuan Geng, Weihua Zhao, Shuxia Yao, Debo Dong, Bharat B. Biswal, Keith M. Kendrick, Benjamin Becker

## Abstract

Attention and salience processing have been linked to the intrinsic between- and within-network dynamics of large scale networks engaged in internal (default mode network, DN) and external attention allocation (dorsal attention, DAN, salience network, SN). The central oxytocin (OXT) system appears ideally organized to modulate widely distributed neural systems and to regulate the switch between internal attention and salient stimuli in the environment. The current randomized placebo (PLC) controlled between-subject pharmacological resting-state fMRI study in N = 187 (OXT, n = 94; n = 93; single-dose intranasal administration) healthy male and female participants employed an independent component analysis (ICA) approach to determine the modulatory effects of OXT on the within- and between-network dynamics of the DAN-SN-DN triple network system. OXT increased the functional integration between subsystems within SN and DN and increased functional segregation of the DN with the SN and DAN engaged in attentional control. Whereas no sex differences were observed, OXT effects on the DN-SN interaction were modulated by autism traits. Together, the findings suggest that OXT may facilitate efficient attentional allocation towards social cues by modulating the intrinsic functional dynamics between DN components engaged in social processing and large-scale networks involved in external attentional demands (SN, DAN).

## Introduction

Attention represents a cardinal, yet strongly limited, cognitive resource. The capacity to flexibly switch between internally focused attention and external attentional demands, including both top-down orientation in response to task demands as well as bottom-up reorientation towards salient stimuli in the environment, represents a key mechanism promoting efficient allocation of this limited resource (Buschman and Miller, 2007; Corbetta et al., 2008). Accumulating evidence suggests that the interplay between competing attentional processes is mirrored in the intrinsic functional organization of the brain, particularly the temporal anti-correlations between the default network (DN), engaged during rest and internally-oriented processes (Buckner et al., 2008; Raichle et al., 2001) and networks engaged during external attentional demands such as the dorsal attention (DAN) and salience networks (SN) (Corbetta and Shulman, 2002; Fox et al., 2006; Fox et al., 2005; Menon and Uddin, 2010; Poole et al., 2016).

The DN comprises a set of interacting subsystems centered at the precuneus/posterior cingulate cortex (PCC), anterior medial prefrontal cortex (mPFC), medial temporal and inferior parietal regions that are consistently suppressed during externally-oriented attention, yet more active during internally-oriented and self-referential processes (Andrews-Hanna et al., 2010; Buckner et al., 2008; Raichle et al., 2001). In contrast, the DAN and SN support externally-oriented attentional processes, with the DAN centered at the frontal eye field (FEF) and anterior intraparietal sulcus extending into the superior parietal lobule (aIPS/SPL) specifically contributing to goal-directed top-down attention (Christoff et al., 2016; Corbetta and Shulman, 2002; Fox et al., 2006). On the other hand, the SN centered at the anterior insula (AI) and dorsal anterior cingulate cortex (dACC), plays an important role in the bottom-up detection of salient external stimuli and stimulus-guided attentional control (Corbetta et al., 2008; Menon and Uddin, 2010; Seeley et al., 2007).

The DAN and DN have been characterized by their competitive relationship during both rest and external goal-directed attentional control (Dixon et al., 2017; Fox et al., 2005; Gao and Lin, 2012; Kelly et al., 2008; Spreng et al., 2010). The behavioral relevance of the intrinsic anti-correlation is further supported by associations with individual differences in goal-directed task performance (Kelly et al., 2008). Furthermore, accumulating evidence suggests that SN and DN operate in a competitive relationship, and that in particular the right AI represents a critical node for suppressing DN activity and reallocating attentional resources in response to external salient events (Andrews-Hanna et al., 2014; Menon and Uddin, 2010; Uddin, 2015). The SN has additionally been proposed to mediate internally and externally directed attention by maintaining a dynamic balance between the DAN and DN (Smallwood et al., 2012; Spreng et al., 2010). Disruptions in the dynamic interplay between these networks, particularly attenuated anti-correlations with the DN, have been observed across mental disorders characterized by marked deficits in attention and salience processing, including attention deficit hyperactivity, autism spectrum and psychotic disorders (Anticevic et al., 2012; Castellanos and Aoki, 2016; Dong et al., 2018; Ebisch and Aleman, 2016; Padmanabhan et al., 2017; Sidlauskaite et al., 2016; Zhang and Raichle, 2010). From a conceptual perspective, the dynamic interplay between the DN, SN and DAN has been considered as a key intrinsic organizational principle of the brain that may facilitate functional integration across segregated networks (Zhou et al., 2017) and efficient attentional control (Corbetta and Shulman, 2002; Fox et al., 2006; Fox et al., 2005; Menon and Uddin, 2010; Poole et al., 2016). For instance, Poole et al. (2016) found that the intrinsic functional connectivity between and within the competing DN, DAN and SN predicts individual differences in distractor suppression and cognitive performance.

By combining resting-state fMRI (rsfMRI) with pharmacological challenges research has recently begun to explore how different neurotransmitter systems modulate the intrinsic functional organization of large-scale brain networks (overview in Khalili-Mahani et al., 2017; Lu and Stein, 2014). With respect to the functional dynamic architecture of the attention-related DAN-SN-DN system, the hypothalamic neuropeptide oxytocin (OXT) appears of particular interest. OXT regulates functional domains that have been strongly associated with the DAN-SN-DN systems, including not only attention and salience processing but also social cognition, interoception and self-referential processes (Hurlemann and Scheele, 2016; Johnson and Young, 2017; Ma et al., 2016; Meyer-Lindenberg et al., 2011; Shamay-Tsoory and Abu-Akel, 2016). During task challenges, modulatory effects of OXT on regional activity within the DAN-SN-DN networks have been consistently reported (Wigton et al., 2015), with accumulating evidence suggesting that effects in DN regions may mediate OXT’s influence on self-referential and social-feedback processing (Baumgartner et al., 2008; Eckstein et al., 2014; Hu et al., 2015; Labuschagne et al., 2012; Liu et al., 2017; Zhao et al., 2017; 2016). On the other hand, effects of OXT on salient external cues (Luo et al., 2017; Riem et al., 2011) and reorienting attention towards them (Yao et al., 2017) are mediated by modulation of SN regions. In line with the role of the DAN in directed attention, OXT enhanced activity in DAN core nodes during explicit processing of social stimuli (Levy et al., 2016) and active engagement in social interactions (Baumgartner et al., 2008; Rilling et al., 2012).

Furthermore, from a physiological perspective the central OXT system appears ideally organized to modulate widely distributed neural systems (Bethlehem et al., 2013; Johnson and Young, 2017; Mitre et al., 2017). Relevant properties of the OXT system include not only its central projection profile, somatodendritic release and widely distributed expression of OXT-sensitive receptors in the brain, but also a long half-life after release (20 min) as well as interactions with classical neurotransmitter such as serotonin and dopamine (Bethlehem et al., 2013; Johnson and Young, 2017; Mitre et al., 2017). In line with this conceptual framework, a growing number of recent studies have reported modulatory effects of OXT on the intrinsic functional interplay between widely distributed brain regions across species (e.g. Bethlehem et al., 2017; Eckstein et al., 2017; Rubin et al., 2017; Wang et al., 2017).

Against this background, the present randomized, between-subject, placebo-controlled pharmacological fMRI study in a large sample of healthy male and female participants (*N* = 187) aimed at determining modulatory effects of OXT on the intrinsic organization of, and the dynamic interplay between, the DAN-SN-DN systems. In the context of increasing evidence for sex-differential effects of intranasal OXT on neural activity during externally-oriented social information processing (Gao et al., 2016; Luo et al., 2017; Rilling James et al., 2018), we additionally aimed to determine whether sex-differential effects can be additionally observed at the level of intrinsic brain functional connectivity. Finally, a disrupted interplay between the DAN-SN-DN has been consistently observed in autism (e.g. Uddin et al., 2017) and considerable efforts aim at evaluating the therapeutic potential of intranasal OXT for alleviating symptoms of autism (Guastella and Hickie, 2016; Keech et al., 2018). Thus, in line with previous studies (Bethlehem et al., 2017; Scheele et al., 2014) associations between OXT effects and individual variations in autistic traits were additionally explored. To delineate the functional interaction between SN, DAN and DN on a finer scale, the large-scale networks were initially determined at the subsystem-level, next the effects of OXT on intrinsic connectivity within and between these subsystems were determined. Based on the physiological properties of the OXT system, we expected that OXT would particularly modulate between-network interactions while sparing within-network organizations. Furthermore, in line with previous research we expected that the effects of OXT would vary as a function of individual levels of autistic traits.

## Materials and Methods

### Participants

Figure 1 provides an overview of the experimental protocols and analysis pipeline. *N* = 197 healthy, right-handed university students were enrolled in the study. *N* = 10 participants were excluded due to excessive head motion (>2.5 mm translation, >2.5° rotation) leading to a final sample size of *N* = 187 (randomized double-blind allocation to 94 = OXT, 93 = placebo-administration (PLC); mean age = 21.4 ± 2.15 years; 50.8% males). All participants were free from current or past psychiatric, neurological or other medical disorders. Participants were instructed to abstain from alcohol and caffeine during the 24 hours prior to the experiment. All female participants were nulliparous and none was pregnant or using hormonal contraceptives. χ^2^ test revealed no significant differences between the treatment groups with respect to menstrual cycle phase (follicular, luteal phases, *P* = 0.832, two-sided). Written informed consent was obtained, study procedures had full ethical approval by the local ethics committee and were in accordance with the latest revision of the declaration of Helsinki.

**Figure 1.**
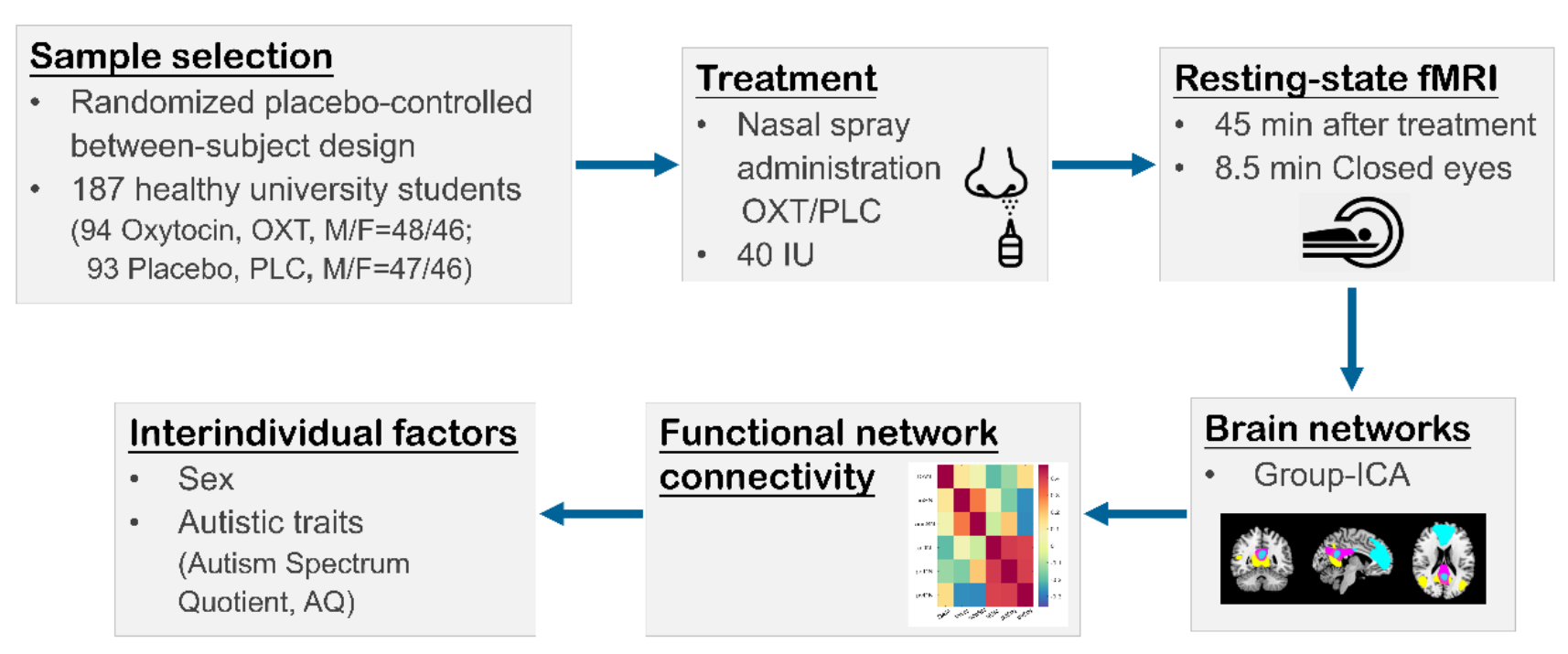
Experimental procedures and key processing steps.

### Psychometric assessment

Before drug administration, participants completed the State-Trait Anxiety Inventory (STAI) (Spielberger, 1983) and the Positive and Negative Affect Schedule (PANAS) (Watson et al., 1988) scales to control for effects of current emotional state on the resting state activity. The Beck’s Depression Inventory (BDI) (Beck et al., 1996) was administered to control for confounding effects of different levels of depression between treatment groups. In addition, the Autism Spectrum Quotient (AQ) (Baron-Cohen et al., 2001) scale was administered to assess levels of autistic traits.

### Experimental procedure

In a double-blind randomized placebo-controlled between-subject pharmaco-fMRI design, participants self-administered either a single dose of 40 international units (IU) OXT (ingredients: oxytocin, glycerin, sodium chloride and purified water; Sichuan Meike Pharmaceutical Co. Ltd, Sichuan, China) or placebo (PLC) nasal spray (produced by Sichuan Meike Pharmaceutical Co. with same ingredients except for OXT) corresponding to 5 puffs per nostril. The dosage was chosen in line with previous OXT studies (Frijling et al., 2016; Geng et al., 2018; Koch et al., 2016; Shin et al., 2018; Yao et al., 2017; Zhao et al., 2017), and adminstered in accordance with guidelines for OXT administration in humans (Guastella et al., 2013). We and other groups have shown that this 40 IU dose produces either similar or improved behavioral and neural effects compared to a 24 IU dose in task-related paradigms (Geng et al., 2018; Shin et al., 2018; Zhao et al., 2017). Based on pharmaco-kinetic experiments (Paloyelis et al., 2016) treatment was administered 45 min before acquisition of the functional MRI time-series.

### fMRI acquisition

MRI data was collected on a 3-Tesla GE MR750 Discovery MRI system (General Electric Medical System, Milwaukee, WI, USA). High-resolution brain structural data was acquired using a T1-weighted sequence (repetition time (TR) = 5.97 ms, echo time (TE) = 1.97 ms, flip angle = 9°, field of view (FOV) = 256 × 256 mm, image matrix = 256 × 256, slice thickness = 1 mm, and 128 sagittal slices) to improve spatial normalization of the functional data. A total of 255 volumes were acquired using a T2*-weighted Echo Planar Imaging (EPI) sequence with the following parameters: TR = 2000 ms, TE = 30 ms, FOV = 240 × 240 mm, flip angle = 90°, image matrix = 64 × 64, thickness/gap =3.4/0.6 mm, 39 axial slices with an interleaved ascending order. During the 8.5 min resting-state scanning, the participants were instructed to lay still while keeping their eyes closed and relax to let the mind wander while not falling asleep. Head movements were minimized by using comfortable head cushions. In post scan interviews none of the participants reported having fallen asleep.

### fMRI data analysis

#### Functional network definition

Data preprocessing was carried out using SPM12 (http://www.fil.ion.ucl.ac.uk/spm/, Wellcome Trust Centre for Neuroimaging). In order to allow for MRI signal equilibrium, the first 10 volumes were removed. The preprocessing steps included slice timing, head motion correction, spatial normalization and smoothing (8 mm full-width at half maximum Gaussian kernel). Subject head motion was assessed using mean framewise displacement (FD) (Power et al., 2012) and importantly there was no significant difference in mean FD between the treatment groups (OXT: Mean ± SD = 0.097 ± 0.036; PLC: Mean ± SD = 0.096 ± 0.043; *P* = 0.822, two-sample t-test).

Next, group independent component analysis (ICA) was performed using GIFT (http://icatb.sourceforge.net/) (Calhoun et al., 2001) to retrieve large-scale networks and identify our networks of interest. In line with recent recommendations, a data-driven parcellation (ICA) was employed to increase the accuracy for determining the networks in the dataset that is used for subsequent analyses (in contrast to an atlas-based approach) (Bijsterbosch et al., 2017). To prevent a biased determination of the ICA components, these were initially defined using the pooled data from both groups (Bijsterbosch et al., 2017). The optimal number of components was set to 25 based on previous work (Shirer et al., 2012; Tsvetanov et al., 2016; Xin and Lei, 2015) demonstrating that this number can reliably determine the major large-scale intrinsic networks and their subsystems, including the DAN, SN and DN that were of particular interest for the present research question. To ensure the robustness of the estimated Independent Components (ICs), the Infomax ICA algorithm (Bell and Sejnowski, 1995) was repeated 10 times using ICASSO (Himberg et al., 2004). ICs and time courses for each participant were back-reconstructed, and the mean spatial maps were transformed to z-scores for display purposes. Using a network template approach based on the Stanford Resting State Network templates (http://findlab.stanford.edu/functional_ROIs.html) (Shirer et al., 2012), six ICs of interest were determined: DAN, anterior cingulate salience network (accSN), insula salience network (inSN), anterior default network (aDN), posterior dorsal default network (pdDN) and posterior ventral default network (pvDN). The spatial distribution of six ICs is presented in Figure 2, and the peak coordinates of the group-level regions within each network are summarized in Table 1. Additional processing steps were applied to the time courses of ICs of interest to improve noise control, including detrending (linear, cubic and quadratic), regression of motion parameters and their temporal derivatives, despiking detected outliers, and low-pass filtering using a high-frequency cutoff at 0.15 Hz (Allen et al., 2014).

**Figure 2.**
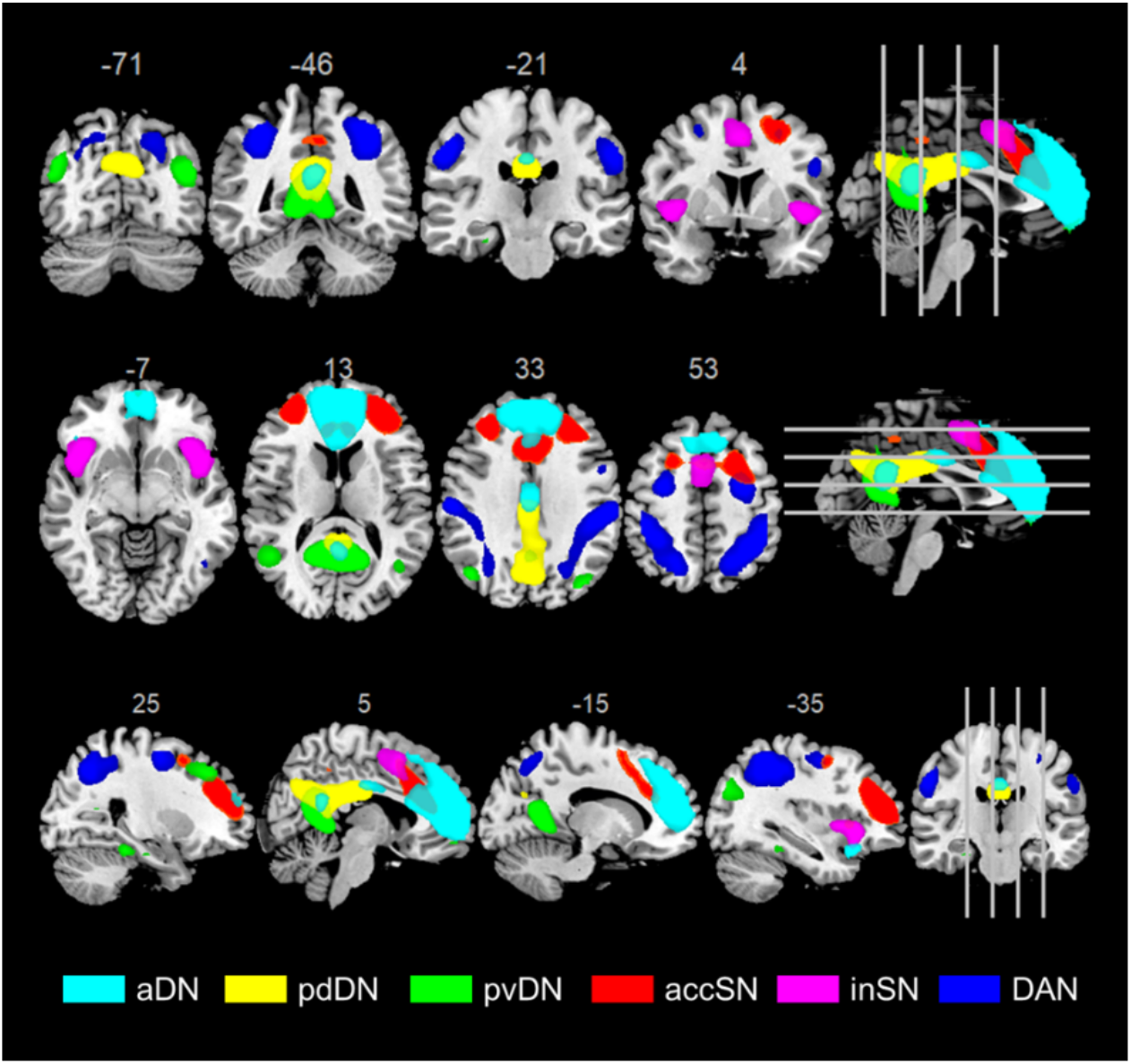
Spatial distribution of the identified large-scale network components. Abbreviations: aDN, anterior default network; pdDN, posterior dorsal default network; pvDN, posterior ventral default network; accSN, anterior cingulate salience network; inSN, insula salience network; DAN, dorsal attention network.

**Table 1.**
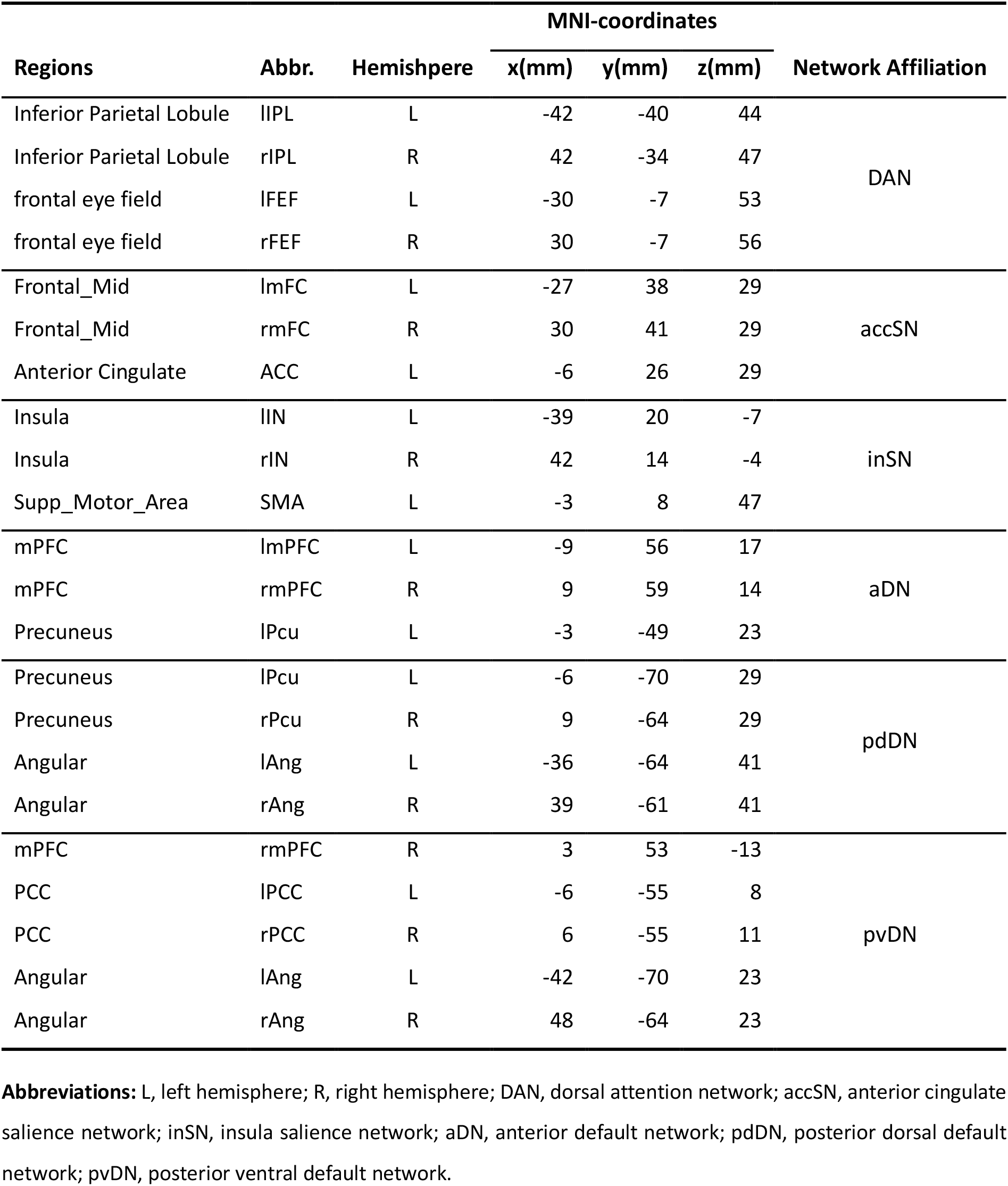
Peak foci on the group-level for the DAN, accSN, inSN, aDN, pdDN and pvDN defined by ICA in the pooled sample (OXT and PLC).

#### Meta-analytic decoding of network function using NeuroSynth

The functional properties of each network were decoded using a large-scale database-informed meta-analytic approach as implemented in NeuroSynth (Yarkoni et al., 2011). The 16 terms showing the highest correlations for each subsystem mask were extracted, and the terms corresponding to each subsystem visualized as a word cloud (Figure 3). The font size of the term in each word cloud is proportional to the correlation strength.

**Figure 3.**
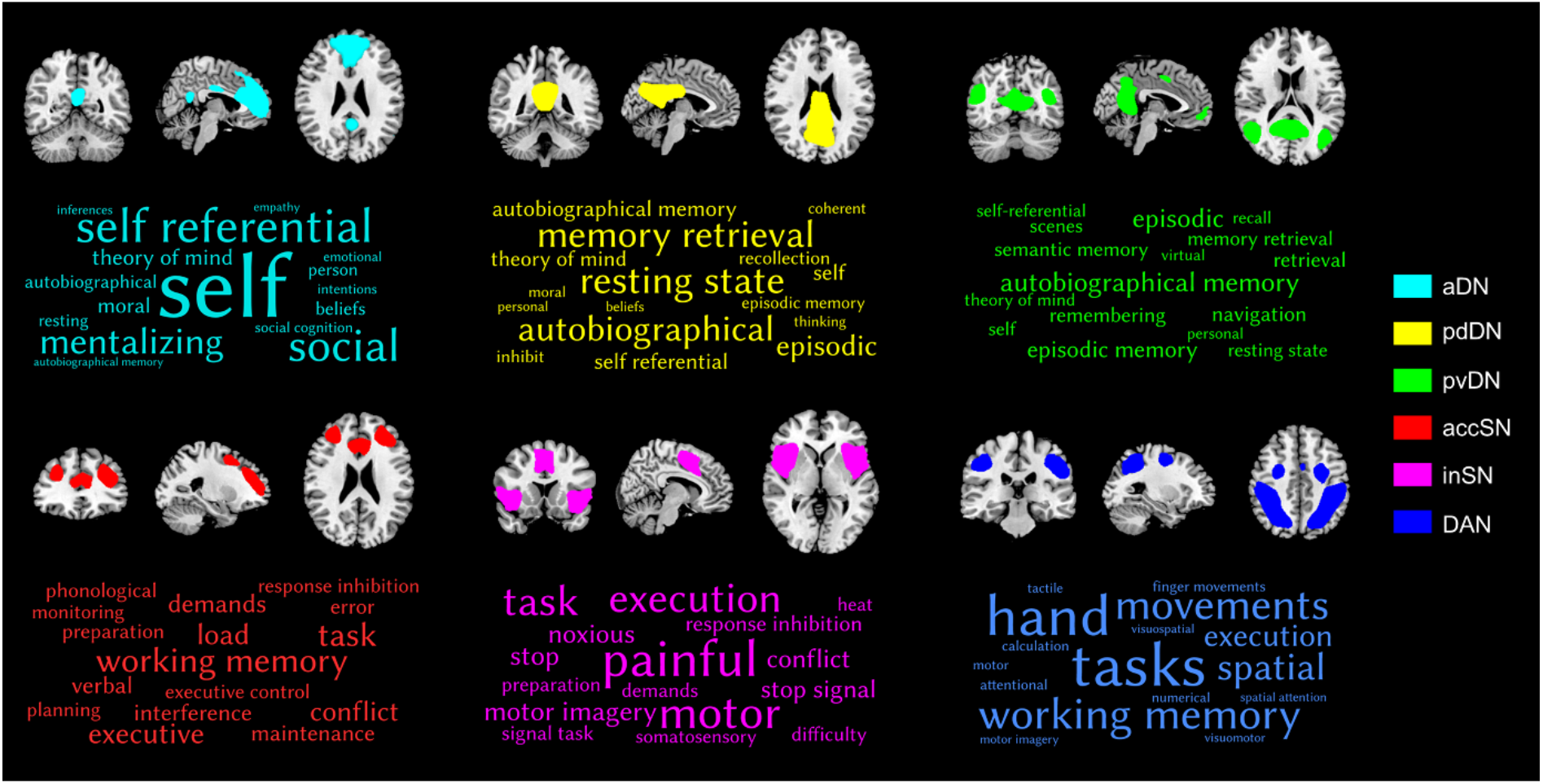
Decoding the functional properties of the identified networks using NeuroSynth. Only the top 16 terms are visualized. The font size reflects the size of the correlation. Abbreviations: aDN, anterior default network; pdDN, posterior dorsal default network; pvDN, posterior ventral default network; accSN, anterior cingulate salience network; inSN, insula salience network; DAN, dorsal attention network.

#### Functional network connectivity and statistical analysis

##### Between-network connectivity analysis

Pearson correlations between the time courses of DAN, accSN, inSN, aDN, pdDN and pvDN components were calculated to determine between-network functional connectivity. The correlation coefficients were normalized to z-scores with Fisher’s r-to-z transformation to increase the normality of the distribution, allowing further statistical analysis of correlation strengths. To examine interactions between treatment, sex, and between-network connectivity, correlation coefficients for each participant were subjected to a mixed-effect analysis of variance (ANOVA), with two between-subject factors treatment (OXT, PLC) and sex (males, females) and the within-subject factor network pairs (15 pairs). To further disentangle the direction of effects involving treatment, post hoc comparisons were conducted using independent sample t-tests (two-tailed).

##### Within-network connectivity analysis

The spatial map from group ICA reflect the component’s within-network connectivity pattern (i.e. the mixing weights) in a voxel-wise manner. Voxels highly correlated with the component’s time course receive high connectivity values. To examine voxel-wise level treatment group differences, corresponding spatial z-maps of each network were entered into two-sample t-tests (two-tailed). For the network level analysis of within-network dynamics, subject-specific median incorporating voxels within the respective component were computed for each participant (Myers et al., 2014; Schlaffke et al., 2017). The network median values were subjected to a mixed-effect analysis of variance (ANOVA), with the between-subject factors treatment (OXT, PLC) and sex (males, females) and the within-subject factor network (6 networks).

The false discovery rate (FDR) (Benjamini and Hochberg, 1995) correction was employed when performing multiple statistical tests simultaneously in all the functional networks. The multiple comparison correction threshold is *P*_FDR_ = 0.05. Finally, we used the Pearson correlations to assess the correspondence between the functional network connectivity and the autistic traits scores across participants. Between-group correlation differences were tested using Fisher’s z tests. Based on findings from a previous rsfMRI study reporting stronger OXT network-level effects in subjects with higher autistic traits, between-group correlation differences were tested one-tailed (Bethlehem et al., 2017).

## Results

### Demographics and questionnaires

OXT (*N* = 94) and PLC (*N* = 93) groups did not differ in gender, age, levels of depression, mood, anxiety and autistic traits (detailed group characteristics are given in Table 2).

**Table 2.** Demographics and questionnaires.

| | PLC<br>(N = 93) | OXT<br>(N = 94) | *T*/$\chi^2$ | *P* |
| --- | --- | --- | --- | --- |
| **Age (years)** | 21.39 ± 2.24 | 21.38 ± 2.06 | 0.01 | 0.99 |
| **Gender(M/F)** | 47/46 | 48/46 | 0.005 | 0.94 |
| **STAI-state** | 38.61 ± 7.88 | 38.70 ± 7.32 | -0.08 | 0.94 |
| **STAI-trait** | 41.37 ± 6.91 | 41.23 ± 7.38 | 0.13 | 0.90 |
| **BDI** | 7.14 ± 5.61 | 6.43 ± 5.96 | 0.84 | 0.40 |
| **PANAS-positive** | 29.10 ± 5.92 | 27.88 ± 5.47 | 1.46 | 0.15 |
| **PANAS-negative** | 19.12 ± 6.26 | 17.91 ± 5.73 | 1.37 | 0.17 |
| **AQ** | 20.32 ± 4.97 | 19.36 ± 5.31 | 1.28 | 0.20 |

**Abbreviations:** OXT, oxytocin; PLC, placebo; M, male; F, female; STAI, State-Trait Anxiety Inventory; BDI, Beck's Depression Inventory; PANAS, Positive and Negative Affect Schedule; AQ, Autism Spectrum Quotient. Data are expressed as mean ± SD (SD: standard deviation). For continuous variables, independent sample t-tests were carried out. For categorical variables, $\chi^2$ tests were employed.

### Effects of OXT on between-network connectivity

We performed a three-way repeated-measures ANOVA, with two between-subject factors treatment (OXT, PLC) and sex (males, females) and a within-subject factor network pairs (15 pairs). A significant main effect of network pairs [F(14, 170) = 266.05, *P* < 0.001, η^2^ *P* = 0.956] and sex (F(1, 183) = 9.50, *P* = 0. 002, η^2^ *P* = 0.049) were observed. Moreover, there was a significant interaction between network pairs and sex [F(14, 170) = 47.55, *P* < 0.001] and between network pairs and treatment [F(14, 170) = 23.50, *P* = 0.005]. We did not find either a two-way interaction between sex and treatment (*P* = 0.279) or a three-way interaction between the factors (*P* = 0.649), suggesting that OXT produced comparable effects in male and female participants. Post hoc tests revealed that OXT significantly increased positive connectivity in network pairs: DAN-accSN (t(185) = 2.06, *P* = 0.041, Cohen’s d = 0.302); accSN-inSN (t(185) = 2.01, *P* = 0.046, Cohen’s *d* = 0.294); aDN-pvDN (t(185) = 2.16, *P* = 0.032, Cohen’s *d* = 0.316); pdDN-pvDN (t(185) = 2.77, *P*_FDR_ = 0.031, Cohen’s *d* = 0.405) (Figure 5A). Whereas, OXT significantly increased anti-correlation in network pairs: DAN-aDN (t(185) = −2.52, *P*_FDR_ = 0.047, Cohen’s *d* = 0.369); inSN-pvDN (t(185) = −2.15, *P* = 0.033, Cohen’s *d* = 0.314) (Figure 5B). Finally, OXT significantly reduced the connectivity between network pairs: DAN-pvDN (t(185) = −3.01, *P*_FDR_ = 0.022, Cohen’s *d* = 0.440); accSN-pvDN (t(185) = 3.06, *P*_FDR_ = 0.022, Cohen’s *d* = 0.447) (Figure 5B). Between-network connectivity visualized by means of connectivity matrices, and the modulatory effects of OXT, are schematically summarized in Figure 4. After multiple comparisons using FDR, group differences were only significant in DAN-aDN, DAN-pvDN, accSN-pvDN and pdDN-pvDN (Figure 4 and Figure 5).

**Figure 4.**
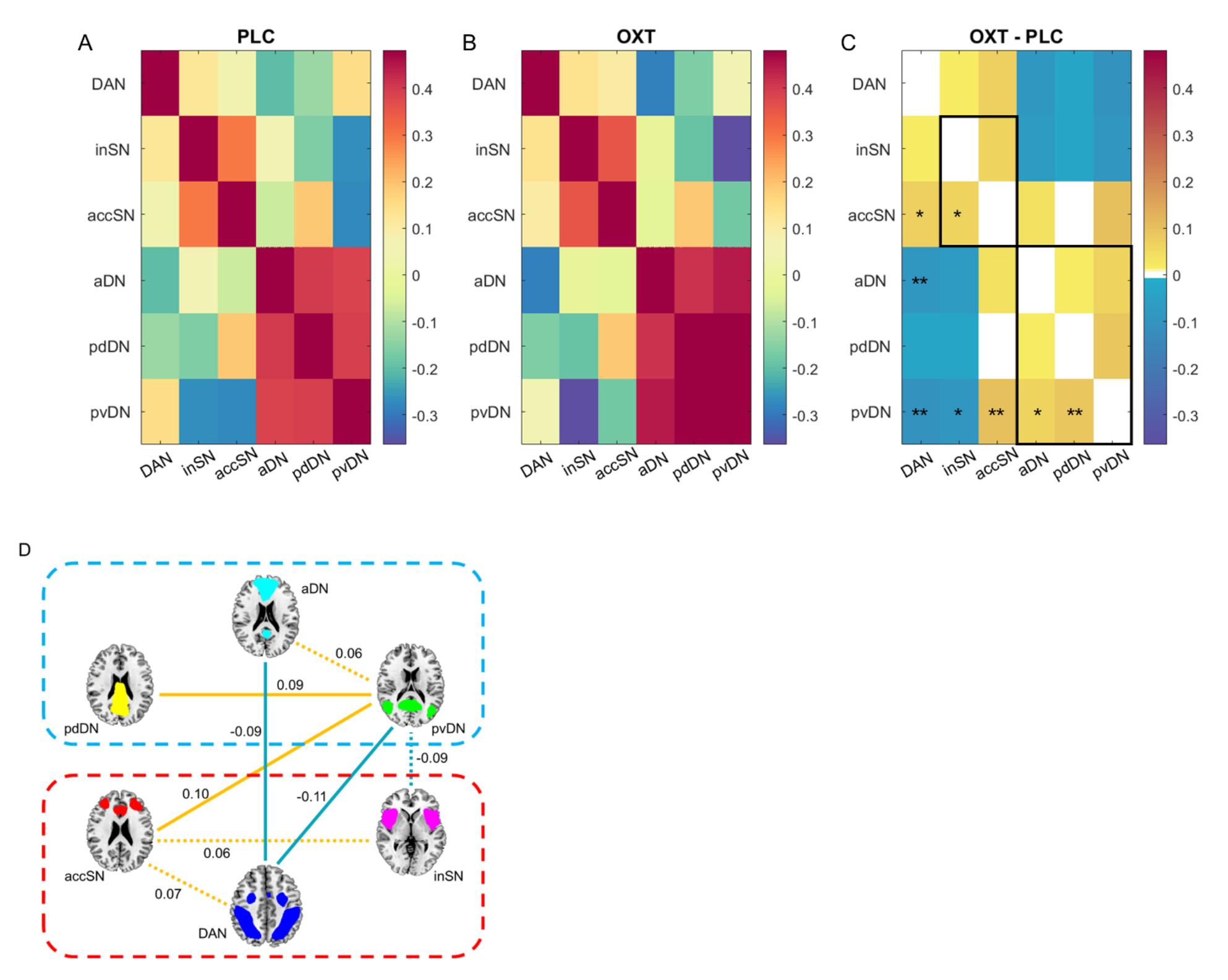
Between-network connectivity for (A) PLC and (B) OXT groups. (C) Group differences (OXT - PLC) in between-network connectivity. * indicates significant differences at *P* < 0.05, uncorrected; ** indicates significant differences at *P* < 0.05, FDR-corrected. (D) A schematic summarizing the modulatory effects of OXT on between-network connectivity. Orange lines indicate increased connectivity; Cyan lines indicate decreased connectivity; Dashed lines indicate significant differences at *P* < 0.05, uncorrected; Solid lines indicate significant differences at *P* < 0.05, FDR-corrected. Abbreviations: DAN, dorsal attention network; accSN, anterior cingulate salience network; inSN, insula salience network; aDN, anterior default network; pdDN, posterior dorsal default network; pvDN, posterior ventral default network.

**Figure 5.**
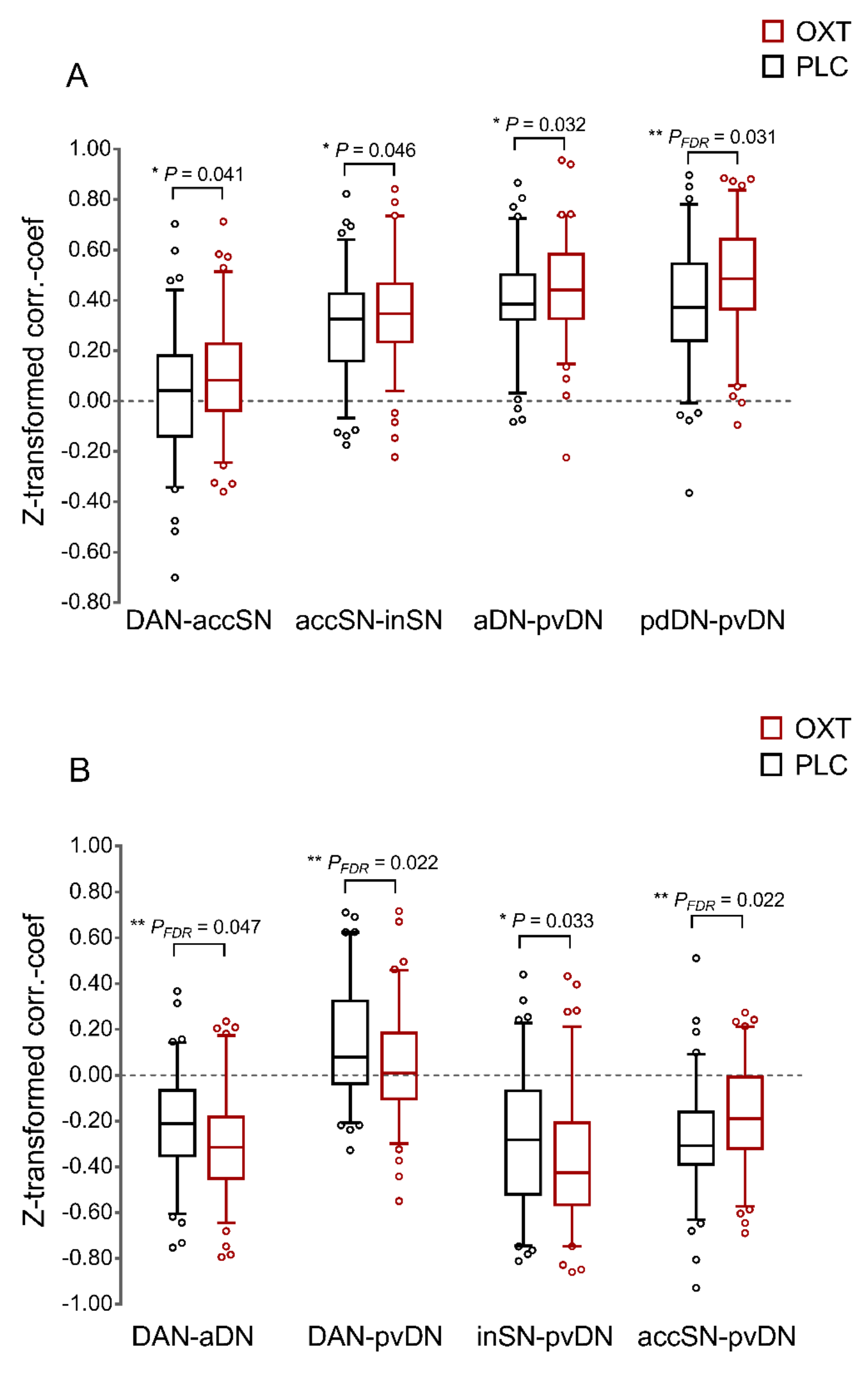
Box plots for significant differences in between-network connectivity. (A) Increasing functional integration after OXT treatment. (B) Increasing functional segmentation after OXT treatment. * indicates significant differences at *P* < 0.05, uncorrected; ** indicates significant differences at *P* < 0.05, FDR-corrected.

### Effects of OXT on within-network connectivity

On both voxel-wise and whole-network levels of the spatial maps, there were no significant differences between the treatment groups with respect to within-network connectivity (all *Ps* > 0.05). Additional ROI-based were carried out to further confirm the lack of effects of treatment on within-network connectivity (see supplementary analyses). Together the results suggest that OXT did not significantly influence within-network organization.

### Associations of between-network connectivity and autism traits

There were no significant differences between treatment groups on AQ scores (all *Ps* > 0.05, see Table 2). We examined whether OXT effects on between-network connectivity were associated with individual variations in autistic traits. Intriguingly, we found some associations between-network connectivity with AQ only in the OXT group. Mean accSN-pvDN connectivity showed significant positive correlation with AQ in the OXT but not in the PLC group (OXT: r(92) = 0.32, *P* = 0.001; PLC: (r(91) = 0.049, *P* = 0.64; Fisher’s *z*: z = 1.931, *P* = 0.027, one-tailed; Figure 6A). Mean accSN-pdDN connectivity showed significant positive correlation with AQ in the OXT but not in the PLC group (OXT: (r(92) = 0.22, *P* = 0.034; PLC: r(91) = −0.042, *P* = 0.69; Fisher’s *z*: z = 1.780, *P* = 0.038, one-tailed; Figure 6B). Mean inSN-pdDN connectivity showed significant positive correlation with AQ in the OXT but not in the PLC group (OXT: (r(92) = 0.27, *P* = 0.009; PLC: r(91) = −0.09, *P* = 0.39; Fisher’s *z*: z = 2.448, *P* = 0.007, one-tailed; Figure 6C). In general, OXT induced a positive association between autistic traits and DN-SN interaction whereas no such association was observed under PLC. We did not observe any significant associations between within-network connectivity and AQ scores (all *Ps* > 0.05).

**Figure 6.**
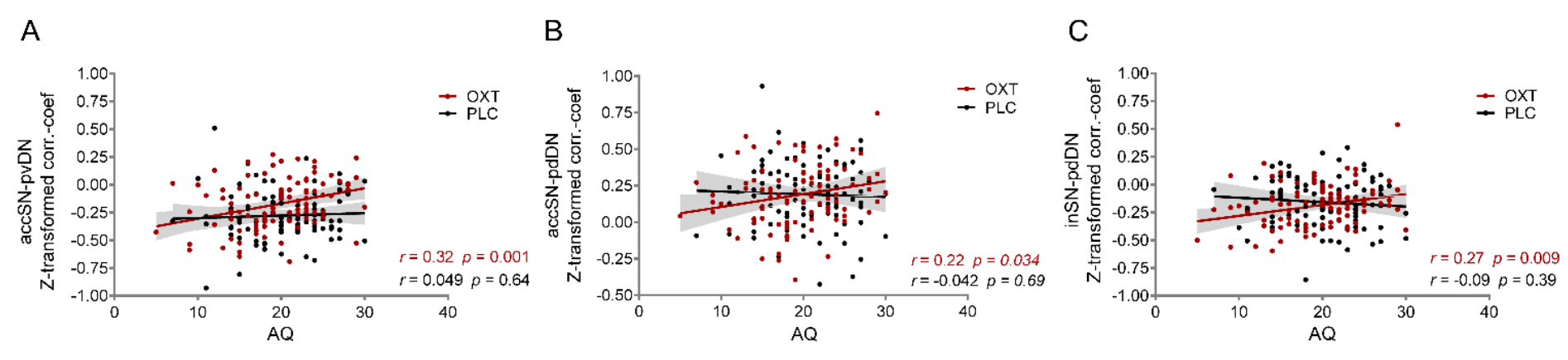
Relationship between AQ scores and between-network connectivity: (A) accSN-pvDN, (B) accSN-pdDN and (C) inSN-pdDN. OXT (red) and PLC (black).

## Discussion

Accumulating evidence suggests that efficient allocation of attentional resources is promoted by the antagonistic dynamics between large-scale brain networks involved in internal information processing (DN) and networks engaged during external attentional demands (DAN, SN) (Fox et al., 2005; Fox et al., 2009; Fransson, 2005; Kelly et al., 2008). In line with previously proposed physiological and behavioral characterizations of the OXT system, the present pharmaco-rsfMRI study revealed that intranasal OXT modulated the between-network dynamics of the DN with the DAN and SN. OXT primarily affected the intrinsic dynamics between the pvDN subsystem with both networks engaged in external attentional demand (SN, DAN) and the dorsal posterior DN, as well as between the anterior DN sub-component and the DAN. Whereas no sex-differences were observed, exploratory analyses revealed that the levels of autistic traits may mediate OXT’s modulatory effects on the intrinsic organization of the attention-related large-scale network architecture.

The ICA approach revealed three DN and two SN subsystems, resembling sub-components of these systems reported in previous studies (e.g. Damoiseaux et al., 2006; Doll et al., 2015; Schlaffke et al., 2017). Subsequent examination of the identified components confirmed that (1) the identified DN components are functionally characterized by internal, while the DAN/SN are characterized by external information processing domains, and that (2) the DN subsystems show distinct anti-correlations with the DAN and SN (as previously reported by Dixon et al., 2017). Notably, the pvDN subsystem showed a positive correlation with the DAN in the present study. Recently, Dixon et al. (2017) demonstrated that DN subsystems exhibit differential functional relationships with the DAN and that – when global signal regression is not employed – a DN subsystem similar to the present pvDN does not exhibit stable anti-correlations with the DAN, whereas the DN core and medial temporal lobe subsystems exhibited modest but reliable DAN anti-correlations.

### Modulatory effects of OXT on attention-related large-scale network architecture

Examining effects of OXT on the intrinsic dynamics of the attention-related networks revealed that OXT increased functional coupling within the DN (aDN-pvDN, pdDN-pvDN) and the SN/DAN (accSN-inSN, DAN-accSN) which might reflect an OXT-induced enhancement of intrinsic integrity in the internal versus external information processing systems. Concomitantly, OXT increased the anti-correlation between the DN with both the DAN (DAN-aDN, DAN-pvDN) and SN (inSN-pvDN). Together, this global pattern may reflect that highly integrated cortical networks exhibit stronger intrinsic coupling, while segregation between anti-correlated networks increases following OXT treatment.

On the sub-component level OXT particularly affected the dynamic interplay of the pvDN subsystem, which encompasses posterior DN core nodes in the medial and lateral parietal regions, including the PCC, bilateral angular gyrus and tempo-parietal junction, as well as the most anterior regions of the DN (i.e. mPFC). These regions have been associated with self-referential processes and personally-relevant autobiographic memory (Amodio and Frith, 2006; Andrews-Hanna et al., 2010; Saxe, 2006), and the identified pvDN sub-component strongly resembles a network consistently engaged across social-cognitive processes (Mars et al., 2012; Schilbach et al., 2008). Following OXT treatment, the interplay of the pvDN with the DAN and the anterior cingulate SN component decreased, whereas competitive interactions with the anterior insula SN component increased. Within the SN, the AI is considered as a key hub for the detection of salient stimuli in the environment (Uddin, 2015) that initiates the deployment of attentional resources, via engaging the ACC and the fronto-parietal control network (FPCN), while concomitantly suppressing the DN (Menon and Uddin, 2010; Sridharan et al., 2008) to guide appropriate responses to salient stimuli. In contrast, the dACC SN component is more prominently engaged in subsequent response selection and as an integral part of the executive control network (Petersen and Posner, 2012; Spreng et al., 2013) governs cognitive resources (Bush, 2011; Menon, 2015). Previous task-based OXT-administration studies suggest that the AI mediates OXT’s effects on salience processing (Striepens et al., 2012) as well as the switch from internally directed attention towards external salient cues (Yao et al., 2017). The pattern of enhanced pvDN-inSN anti-correlations in the context of enhanced correlations within the external attentional control networks (DAN-accSN, acc-inSN) may thus reflect a mechanism that gates OXT’s salience enhancing effects of personally relevant stimuli (Hurlemann and Scheele, 2016; Johnson and Young, 2017; Ma et al., 2016; Meyer-Lindenberg et al., 2011; Shamay-Tsoory and Abu-Akel, 2016).

Whereas OXT increased the anti-correlations of the pvDN with the inSN, it concomitantly enhanced the anti-correlations of the aDN with the DAN. The competitive dynamic between the DN and the DAN has been considered as an adaptive intrinsic organizational principle that subserves effective attention by preventing interference between externally and internally information processing (Fox et al., 2005; Fransson, 2005). Supporting evidence comes from studies reporting that stronger DN-DAN anti-correlations associate with better behavioral performance in functions that critically underlie attentional control, such as distractor suppression and behavioral variability (Kelly et al., 2008; Poole et al., 2016), and those reporting that competitive functional organization is attenuated in populations with marked impairments in externally-oriented attention (Buckner et al., 2008; Spreng et al., 2016; Zhang and Raichle, 2010). The ACC and adjacent mPFC integrate emotionally salient signals from limbic and striatal regions to guide attention (Peters et al., 2016), and contribute to self-referential and mentalizing processes (Schmitz and Johnson, 2007; Somerville et al., 2010). In line with the contributions of the mPFC to these functional domains, task-based OXT-administration studies reported that the effects of intranasal OXT on social-cognitive and self-referential processes are mediated by mPFC regions that map onto the aDN (Baumgartner et al., 2008; Labuschagne et al., 2012; Liu et al., 2017; Petrovic et al., 2008; Zhao et al., 2017; 2016).

Together, the enhanced anti-correlations of the DN with the SN and DAN may reflect OXT-induced modulations of the intrinsic attention-network architecture that gate its subsequent salience and attentional processing effects in interaction with environmental stimuli. However, previous animal and human research did not reveal cognitive enhancing properties of OXT per se (Tollenaar et al., 2017; Wirth, 2015), but rather emphasize that OXT specifically facilitates salience and attentional processing of emotional, particularly social, salient stimuli (Harari-Dahan and Bernstein, 2014; Shamay-Tsoory and Abu-Akel, 2016), thereby promoting increased recruitment of attentional resources (Prehn et al., 2013) and processing accuracy for these stimuli (Shahrestani et al., 2013). Importantly, the pvDN and aDN nodes exhibiting OXT-enhanced competitive dynamics with the SN and DAN have been strongly engaged in social-cognitive and self-referential processes (Mars et al., 2012; Schilbach et al., 2008), suggesting that the specific effects of OXT on these sub-components may gate the preferential processing of social and self-relevant stimuli in the environment.

### Sex-dependent effects of OXT

In contrast to an increasing number of task-based OXT administration studies, no sex-differential effects of OXT were observed during the task-free state. Previous studies reported differential effects of OXT on men and women during social evaluation and social interaction (Borland et al., 2018; Feng et al., 2015; Gao et al., 2016) even when stimuli are presented subliminally (Luo et al., 2017). The lack of sex-differential effects during the task-free state suggests that sex-differential effects of OXT evolve in interaction with the social context.

### Association between DN-SN connectivity and autism traits

In line with prior work reporting that OXT’s effect on intrinsic connectivity (Bethlehem et al., 2017) and social processing related neural activity (Scheele et al., 2014) vary as a function of autistic traits, exploratory analysis revealed positive associations between levels of autistic traits and SN-DN (i.e., accSN-pvDN, accSN-pdDN, inSN-pdDN) connectivity following OXT, but not PLC. Both, ACC and AI core regions of the SN have repeatedly been associated with ASD (Anderson et al., 2011; Uddin et al., 2013) and overconnectivity between these networks, possibly reflecting deficient segregation (Rudie et al., 2012), has been considered as promising diagnostic and predictive ASD marker (Hogeveen et al., 2018; Uddin et al., 2017). Initial clinical trials in ASD demonstrated attenuated social-cognitive deficits in ASD following OXT that were mediated by modulatory effects on AI and ACC activity (Andari et al., 2016; Aoki et al., 2014; Watanabe et al., 2015). In this context, OXT-induced the stronger attenuation of accSN-pvDN anti-correlation in participants with higher autistic traits may lend further support for the therapeutic potential of OXT to modulate key neural alterations in ASD.

### Global network perspective

Anti-correlation between task-related networks (i.e., DAN and SN) and the DN is a robust feature of the functional network architecture of the brain (Fox et al., 2005; Fox et al., 2009; Fransson, 2005; Kelly et al., 2008). The antagonism between anatomically and functionally segregated brain systems is a key predictor of efficient behavioral control, and reduced functional segregation between task-related networks and the DN has been proposed as a key factor underlying the imbalance between internal- and external-oriented information processing across mental disorders characterized by impaired attentional control (Cocchi et al., 2013; Mulders et al., 2015; Rudie et al., 2012; Wig, 2017). The proposed competitive intrinsic organization was replicated in the PLC group, with OXT further increasing the anti-correlations of the DN with both attention-related task-networks (SN, DAN). Together with the enhanced positive correlations between DN and SN/DAN sub-components, this may suggest that the OXT system plays an important role in the intrinsic organization of the brain and may promote attentional processing via increasing the functional segregation between attentional networks processing internal and external cues.

## Conclusions

Oxytocin plays an important role as a regulator of attention towards social cues. The current study explored the effect of OXT on intrinsic functional integration and segmentation among the attention-related DAN-SN-DN triple network system. We demonstrated that OXT induced more positive connectivity between subsystems within SN and DN, whereas more DAN-DN and SN-DN anti-correlations with the exception of accSN-DN. Furthermore, we observed an association between SN-DN connectivity and AQ only in OXT group, suggesting OXT’s effect may be more potent for those with higher autistic traits. These results suggest that OXT may facilitate efficient attentional allocation by modulating intrinsic functional interaction between attention-related large-scale brain networks.

## Funding

This work was supported by the National Natural Science Foundation of China (NSFC, 91632117 to BB 31530032 to KMK); the Fundamental Research Funds for Central Universities (ZYGX2015Z002 to BB) and the Science, Innovation and Technology Department of the Sichuan Province (2018JY0001 to BB).

## Disclosure

The authors report no conflict of interests.

